# Interaction between mitochondrial translocator protein and aging in inflammatory responses in mouse hippocampus

**DOI:** 10.1101/2024.06.19.598824

**Authors:** Kei Onn Lai, Nevin Tham, Lauren Fairley, Roshan Ratnakar Naik, Yulan Wang, Sarah R. Langley, Anna M. Barron

**Affiliations:** Lee Kong Chian School of Medicine, Nanyang Technological University Singapore, Singapore; Singapore Phenome Centre, Nanyang Technological University, Singapore 636921, Singapore; School of Biosciences, Cardiff University, Cardiff, United Kingdom

**Keywords:** Aging, hippocampus, mitochondria, neuroinflammation, translocator protein

## Abstract

The mitochondrial translocator protein (TSPO) is a biomarker of inflammation which is upregulated in the brain in aging and associated neurodegenerative diseases, such as Alzheimer’s disease (AD). Here we investigated the interaction between aging and TSPO immunomodulatory function in mouse hippocampus, a region severely affected in AD. Aging resulted in a reversal of TSPO knockout transcriptional signatures following inflammatory insult, with TSPO deletion drastically exacerbating inflammatory transcriptional responses in the aging hippocampus whilst dampening inflammation in the young hippocampus. Drugs that disrupt cell cycle and induce DNA-damage such as heat shock protein and topoisomerase inhibitors were identified to mimic the inflammatory transcriptional signature characterizing TSPO-dependent aging most closely. This TSPO-aging interaction is an important consideration in the interpretation of TSPO-targeted biomarker and therapeutic studies, as well as *in vitro* studies which cannot model the aging brain.

## 1 Introduction

Aging is tightly associated with chronic inflammation (Li *et al*, 2023), and both aging and inflammation are important risk factors for the development of Alzheimer’s disease (AD) (Hou *et al*, 2019; International, 2023; Novoa *et al*, 2022). Key mediators of the inflammatory response in aging and AD are the resident innate immune cells of the brain, microglia. These long-lived cells are susceptible to dysfunction in aging (Conde & Streit, 2006; Hefendehl *et al*, 2014; Lopes *et al*, 2022; Olah *et al*, 2018; Thomas *et al*, 2022) and have been causally implicated in AD pathogenesis by genetic studies (Kunkle *et al*, 2019; Raj *et al*, 2014). Aged microglia exhibit chronic overactivity, including increased proinflammatory cytokine release (Sierra *et al*, 2007; Ye & Johnson, 1999), and dysfunctional immune defences (Hefendehl *et al*., 2014; Stuesse *et al*, 2000; Thomas *et al*., 2022). For example, phagocytosis is impaired in aged microglia, which is an important protective function for mediating clearance of toxic beta amyloid that aggregates in the AD brain (Stuesse *et al*., 2000; Thomas *et al*., 2022). Recently, accelerated inflammatory aging has also been clinically associated with AD (Cullen *et al*, 2021). Consequently, how aging and inflammation, also known as inflammaging, intersect to increase risk of AD is an important question.

The translocator protein, or TSPO, is a useful molecular imaging target to visualize brain inflammaging in health and AD (Bradburn *et al*, 2019; Cagnin *et al*, 2001; Dani *et al*, 2018; Edison *et al*, 2008; Fan *et al*, 2015; Hamelin *et al*, 2016; Kreisl *et al*, 2013; Parbo *et al*, 2017; Schaum *et al*, 2020; Varrone *et al*, 2015; Versijpt *et al*, 2003; Yasuno *et al*, 2012; Yokokura *et al*, 2017). Using positron emission tomography, TSPO signals have been found to significantly correlate with age, AD pathology and cognitive deficits (Dani *et al*., 2018; Edison *et al*., 2008; Fan *et al*., 2015; Finze *et al*, 2023; Hamelin *et al*., 2016; Kreisl *et al*., 2013; Parbo *et al*., 2017; Pascoal *et al*, 2021; Versijpt *et al*., 2003). A mitochondrial protein, TSPO is functionally important in microglial metabolic and immune responses. We and others have shown that genetic deletion of TSPO impairs microglial bioenergetics and phagocytosis, which was linked to dampened acute inflammatory responses in young adult mouse brain and worsened pathogenesis in models of AD (Fairley *et al*, 2023). Beyond inflammation, TSPO has also been functionally implicated in aging, with a recent genetic study showing that TSPO in glia increases longevity in drosophila (Jullian *et al*, 2024). Given mitochondrial and metabolic dysfunction is a universal hallmark of aging (López-Otín *et al*, 2013), and increasing evidence indicates that mitochondrial metabolism plays a key role in coordinating immune processes (Fairley *et al*, 2021), TSPO may provide a interface between aging and inflammation at the mitochondria in microglia. Here we investigated the potential interaction between TSPO immune function and aging in the mouse hippocampus, a region of the brain severely affected in AD.

## 2 Materials and methods

### 2.1 Animals and treatments

Young (3 months old) and aged (20 months old) wild type (WT, C57BL/6) and homozygous global TSPO knock out (TSPO-KO) mice (Barron *et al*, 2018) were used for this study comparison. All experiments were carried out in accordance with the National Advisory Committee for Laboratory Animal Research guidelines and approved by the by the NTU Institutional Animal Care and Use Committee (IACUC# A0384). To induce inflammation, mice were i.p. injected with phosphate buffered saline or 1mg/kg lipopolysaccharide (LPS, L2880, Sigma Aldrich) for 4 days. Brains were harvested 24 hour post-injection. Prior to brain harvesting, mice were anaesthetized through intraperitoneal injection of 120mg/kg body weight of sodium-pentobarbital, and further undergone cardiac perfusion of ice-cold phosphate buffer saline. Hemibrains were then collected and snap frozen using dry ice cooled 2-methylbutane and stored at -80 °C.

### 2.2 RNAseq data generation & analysis

Frozen hippocampi from WT and TSPO-KO mice (n = 4/group, sex-matched within each group) were used to generate the RNAseq data. Of which, our RNAseq dataset from the 3-months old treatment groups was previously analysed and published examining the effect of TSPO-KO at baseline and in inflammatory responses in young adult mouse hippocampus (Fairley *et al*., 2023) were dissected out of hemibrains at -20°C, homogenised immediately in Trizol and total RNA extracted. Chloroform (1:5 vol: vol) was added to the Trizol homogenate then centrifuged for 12000 rpm, 15 min. The aqueous phase was removed, and total RNA extracted using Qiagen RNeasy mini kit column according to the manufacturer’s instruction. RNA quality and purity was checked using the Agilent 2100 Bioanalyzer with Agilent 6000 Nano Kit. Total RNA of the samples had a minimum of RIN 8.3 with an average of RIN 9 and above. OligodT mRNAseq stranded library was prepared using NEBNext Ultra library preparation kit and sequenced using Illumina HiSeqTM.

Quality control of the FASTQ files was performed using FASTQC. Transcripts were quantified and aggregated at the gene level using the Salmon pseudo-aligner 1.9.0 using the *Mus musculus* genome Ensembl release 99 reference. Differential expression analyses were performed using the *DESeq2* (Love *et al*, 2014) 1.36.0 R package and nominal p-values were corrected for multiple testing using FDR. Log_2_ (fold change) (Log2FC) values were shrunk using the *ashr* 2.2-54 R package (Stephens, 2017) and thresholded at Log2FC ≤ 0.5 or Log2FC ≥ 0.5. Volcano plots of RNAseq were constructed using the *ggplot2* (Wickham, 2016) R package 3.5.0 in an R 4.2.3 environment.

For gene set enrichment, the fGSEA R package 1.24.0 (Gennady *et al*, 2021) was used ranking genes by the Wald statistic from the differential gene expression analysis. Enrichments were run against the Reactome collection genesets for *Mus musculus* using 10000 permutations.

### 2.3 RNAseq DEGlist overlap analysis

Differentially expressed genes (DEG) lists generated from DESeq2 are compared against each other for significant non-random associations using the GeneOverlap package (Shen, 2023) (version 1.34.0) from Bioconductor. Each DEG list is filtered for significance and either categorised into gene sets as upregulated (FDR<0.05 and LFC≥0.5) or downregulated (FDR<0.05 and LFC≤-0.5). Fisher’s exact test is performed for each gene set where odds ratio represents strength of non-random association between two gene sets.

### 2.4 Deconvolution methods for the estimate of relative abundance of cell types

Estimation of brain cell type abundance (absolute mode, 100 permutations) was performed using CIBERSORTx (Newman *et al*, 2019). The reference signature was prepared from single cell RNAseq of WT mice hippocampus Zeisel dataset (Zeisel *et al*., 2015). Count matrix of this dataset was CPM (counts per million) transformed using Seurat version 4 package (Hao *et al*, 2024). The bulk mixture of our RNAseq count data was normalized as CPM for the deconvolution (same transformation space as the reference signature) using EdgeR package(Robinson *et al*, 2010). Permutational univariate ANOVA was used to evaluate presence of group differences for each cell type (RVAideMemoire version 0.9-83-7 package), followed by Dunn test (rstatix version 0.7.2 package) with FDR-correction for post-hoc pairwise comparison.

### 2.5 Weighted Gene Co-expression Network

For multiWGCNA (Tommasini & Fogel, 2023), WGCNA network was constructed with soft power threshold of 7, along with the default settings in the other arguments. Before setting as input to the multiWGCNA, the expression matrix was normalized using DESeq2 ‘variance stabilizing transformation’ function. Eigengenes of each module were extracted for plotting. PERMANOVA (permutational multivariate ANOVA, Euclidean method, 9999 permutations) was performed using the expression data for members of each corresponding module through linear model containing main effects, genotype and age, and interaction term genotype:age as testing covariates. PERMANOVA was performed using vegan package version 2.6-4. Following PERMANOVA, post-hoc pairwise comparisons was performed using estimated marginal means (EMM) with the same linear model as PERMANOVA. Estimated marginal means was conducted through emmeans package (Searle *et al*, 1980) version 1.10.0. Module driver genes were identified by high intra-modular connectivity defined as >0.9 Pearson’s correlation with the module Eigenene, PC1 of that specific module. Using module driver genes as query input, protein-protein interactions within the input was retrieved via Network Analyst platform (Zhou *et al*, 2019) and STRING database (Szklarczyk *et al*, 2021) (settings: High confidence PPI >900/1000 score, zero-order direct interactions). The final network was pruned using Prize-collecting Steiner Forest (PCSF) algorithm to isolate high confidence subnetwork consisting of the minimal number of edges.

### 2.6 Transcriptional Motif Analysis

Members of module 5, the module exclusively upregulated in TSPOKO aged group, was used as input for findMotifs.pl function in HOMER (Heinz *et al*, 2010) Motif Analysis. Known enrichment of transcriptional motifs that are either 8 or 10bp, and -2000bp till +500bp from the transcriptional start site (TSS) of the input genes were identified against all known motifs (29786 motifs) within HOMER database. Cognate genes (transcription factors) of significantly enriched (BH<0.05) motifs were then extracted from the expression matrix. These transcription factors were then integrated as a protein-protein interaction (PPI) network with module 5 members using Network Analyst software (Zhou *et al*., 2019) and STRING database (Szklarczyk *et al*., 2021).

### 2.7 Connectivity Map Analysis

The Connectivity Map (CMap) database of drug-induced transcriptional profiles was downloaded from Clue.io (Subramanian *et al*, 2017) (https://clue.io/data/CMap2020#LINCS2020, updated in 2021). This database is made up of over 720,000 drug perturbation signatures, derived from treating more than 33,000 small molecule compounds on various cell lines, including Neural Progenitor Cell (NPC) and glioblastoma (GI1). The top 100 of the differentially expressed genes from the inflammaging comparisons of TSPO-KO were used as inputs, to query the drug CMap database. Differential Expressed Genes were filtered for significance (FDR<0.05) and are either Log2FC ≥0.5 or ≤ -0.5. From within the Library of Integrated Network-Based Cellular Signatures (LINCS) database, compounded-treated signatures derived from the NPC (11,195 signatures) and GI1 (2,117) cell lines were independently queried with the TSPO-KO inflammaging signature.

The sscMap algorithm (Zhang & Gant, 2008) was utilised to compute the connectivity score, due to its superior sensitivity (Lin *et al*, 2020; Tham & Langley, 2022). The connectivity score can range from –1 to +1, where a positive connectivity score indicates a transcriptional phenocopying effect of a drug with the query signature, while a negative connectivity score indicates a transcriptional reversal of the drug and query signatures. From the individual connectivity scores, signatures treated with the same perturbation ID (pert_id) were summarised by computing their mean connectivity score.

The significance of the score is determined by randomly permuting the gene composition of the query signature, 1,000 times, to generate the null distribution scores. A connectivity score is considered significant if the absolute connectivity score exceeds 99% of the absolute scores from the null distribution.

### 2.8 AD risk genes

Human AD risk genes previously identified in genome wide association studies (Gosselin *et al*, 2017) were converted to their mouse orthologs using the BioMart package (Smedley *et al*, 2009) in R. For correlation analysis of AD risk genes for each sample group, unnormalized counts was first transformed using variance stabilizing transformation (blinded to design matrix). Correlation was then performed for each gene-pair using spearman correlation.

### 2.9 Nuclear magnetic resonance spectroscopy (NMR) Metabolomics Analysis

Frozen hemibrains from young and aged WT and TSPO-KO mice (n = 4-5/group) were used for NMR analysis. Brain tissues were weighted and subjected to extraction with pre-cooled methanol:Water (2:1) using homogenizer (Precellys, France). The supernatant was collected after centrifugation (13800 g) and the pellet was subjected to a second extraction using the same procedure. The resultant supernatants were pooled followed by drying using Spin-Vac (Eppendorf, Germany). Extracted samples were resuspended in 550 μL of D_2_O buffer (pH=7.4) containing 0.05% sodium 3-trimethylsilyl (2,2,3,3-^2^H4) propionate (TSP). Here TSP is used as chemical shift reference. NMR spectra were recorded from a Bruker Avance III HD 600 MHz NMR spectrometer (Bruker, Germany) equipped with 5mm BBI 600 MHz Z-Gradient high-resolution probe. One dimensional NMR spectra were recorded using standard first increment of NOSEY pulse sequence (recycle delay-G1-90°-t1-90°-G2-tm-90°-acquisition) with water suppression both during recycle delay (4s) and mixing time (100ms). A total of 256 scans were collected into 64k data points with spectral width of 30 ppm at the temperature of 300k. A 90-degree pulse set ∼11 μs and 4 dummy scan were used.

The spectra were manually phased and corrected for baseline distortion and subsequently imported to Amix (Bruker, Germany, version 3.9.15) and region between 0.5∼10ppm were integrated with 0.003 ppm(1.8Hz) bucket. The water peak resonance (4.60-4.70 ppm) was removed from each spectrum and the spectra were normalized to their respective wet weight of brain tissues. The normalized data were then exported to SIMCA (SIMCA-P15 UMETRICS, Umea, Sweden) for statistical data analysis. Metabolite concentrations were calculated by integration of peak area and further analysed by univariate analysis.

NMR peak assignment was performed by comparison of literature (Abreu *et al*, 2021; Li *et al*, 2015; Wang *et al*, 2005) and some were confirmed with statistical total correlation spectroscopy (STOCSY) (Cloarec *et al*, 2005). A peak at 4.832 ppm was identified that was only present in the aged TSPO-KO mice. Since it is a singlet, two-dimensional NMR is insufficient for the peak assignment. We employed STOCSY method which takes advantage of high correlation for peaks in the same molecule or in the same metabolic pathway (Cloarec *et al*., 2005). It showed high correlation with peak at 8.462, which is formic acid. The correlation suggests that peak at 4.832 could be formaldehyde, which undetectable in normal circumstances. To confirm this assignment, we searched online information. Formaldehyde in water presents structure of methanediol and it is a singlet with chemical shift ranging from 4.4-5.4 ppm (Automated Topology Builder and Repository Version 3.0, Methanediol | CH4O2 | MD Topology | NMR | X-Ray (uq.edu.au)).

Shifted log (log _10_ x+1) transformation was performed on raw data from targeted NMR due to right skewness of the data and presence of multiple zero values. PLSDA (Partial Least Squares Discriminant Analysis) was conducted using MixOmics package version 6.22.0 (Rohart *et al*, 2017). Number of components selected for each corresponding PLSDA was based on the number which gives the least classification error rate for that specified factor (either genotype:age interaction term, main effects genotype or main effects age). Performance of PLSDA was evaluated using 50 repeats and 4-cross fold validation (folds selection based on 4-5 samples per fold). PLSDA plots were generated using MetaboAnalyst 6.0 (Xia *et al*, 2009). Permutational univariate ANOVA (999 permutations) with sample groups as covariate, followed by pairwise post-hoc dunn test was conducted for statistical comparisons. Statistical analysis of the NMR data was performed in an R environment, R 4.2.3 using RVAideMemoire package (Hervé *et al*, 2018) for permutational univariate ANOVA (version 0.9-83-7) and rstatix (Kassambara, 2023) for dunn test (version 0.7.2).

## 3 Results & Discussion

### 3.1 TSPO - aging interaction in inflammatory transcriptional responses in hippocampus

Since TSPO has been implicated in inflammation in brain aging and age-associated neurodegenerative disease, we investigated the role of TSPO in inflammatory responses in the aging mouse hippocampus. We compared the effect of TSPO deletion on LPS-induced transcriptional responses in the hippocampus of aged wild type (WT) and global TSPO knockout (TSPO-KO) mice by RNAseq. TSPO deletion was associated with robust transcriptional differences in the hippocampal inflammatory response in aged mice, with 1184 significant DEGs identified in aged TSPO-KO mice treated with LPS (FDR <0.05; Fig 1A, Supplementary Table 1A). DEGs were further filtered using the shrunken Log2FC of either ≥ 0.5 or ≤ -0.5, resulting in 593 upregulated and 182 downregulated genes (shrunken Log2FC ≥ 0.5 or ≤ -0.5 respectively; Fig 1A, Supplementary Table 1A). Gene set enrichment indicated downregulation of genes belonging to pathways related to the GABAergic synapse, respirasome, ATP synthesis coupled electron transport and NADH dehydrogenase complex assembly and upregulation of genes in pathways involved in immune responses including defense response to symbiont, cell activation involved in immune response, positive regulation of tumor necrosis factor (TNF) superfamily and interleukin (IL) IL-1β production, and response to IL-6 production (Fig. 1B, Supplementary Table 1B). This was surprising since in the young adult mouse hippocampus, we found the opposite effect of TSPO deletion on inflammatory responses, including a dampening of responses to TNF signaling (Fairley *et al*., 2023).

**Figure 1.**
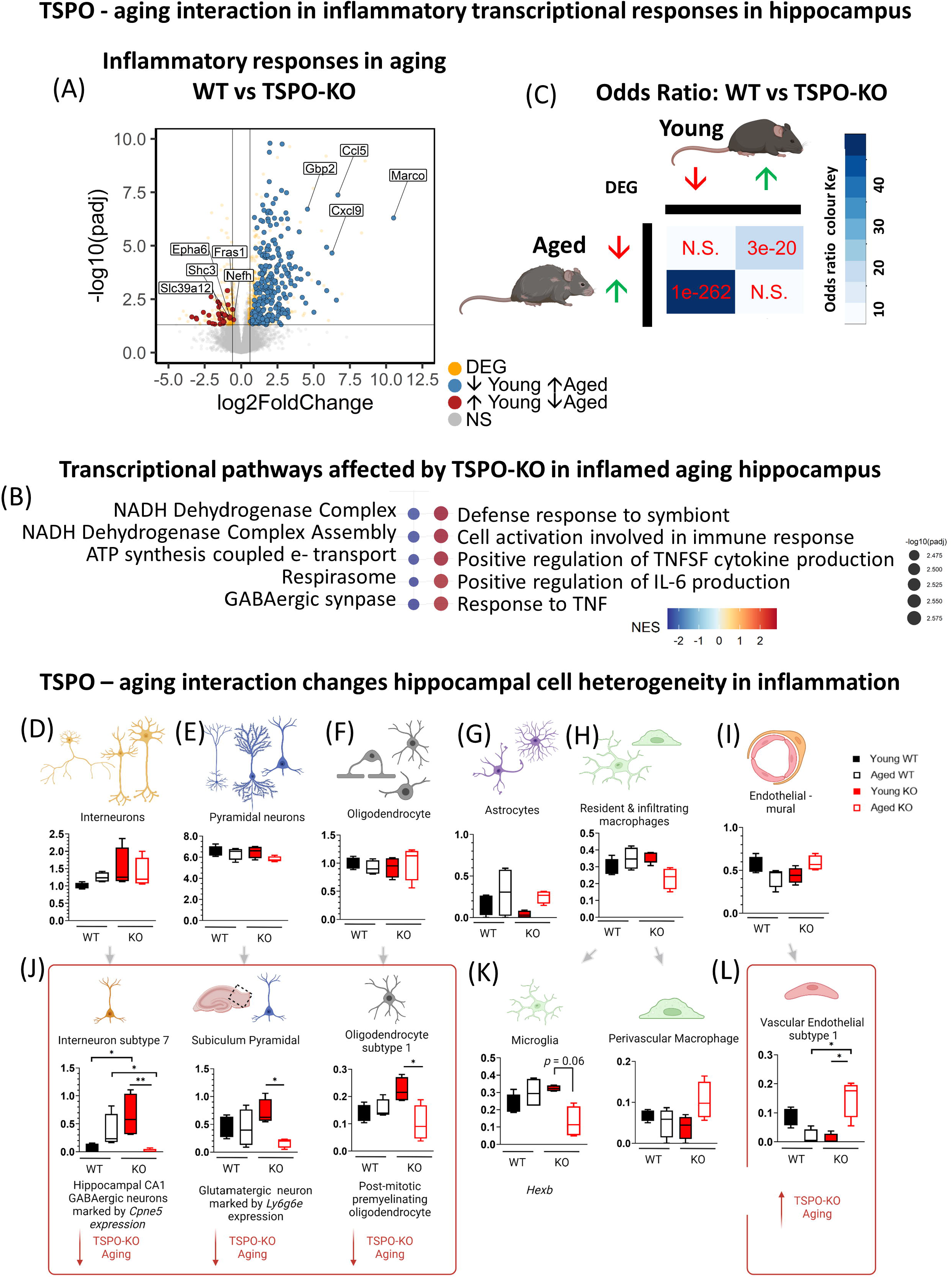
TSPO-aging interaction in inflammatory transcriptional responses in mouse hippocampus. (A) Volcano plot showing differentially expressed genes in aged WT versus TSPO-KO hippocampus following LPS treatment. Differentially expressed genes (DEGs) that were reversed in aged compared to young WT vs TSPO-KO hippocampus indicated in blue (significantly decreased in young but increased in aged) and red (significantly increased in young, but decreased in aged). (B) Fast gene set enrichment analysis (fGSEA) of DEGs in aged WT vs TSPO-KO hippocampus following LPS treatment using Gene Ontology Biological Process pathways. Top 5 up- and down-regulated enriched pathways ranked by normalized enrichment score (NES). (C) Comparison of DEG overlap between aged and young WT vs TSPO-KO comparisons using hypergeometric testing. Odds ratio represents strengths of positive association between two DEG sets. Magnitude of odds ratio is represented by colour key (blue palette) while p-value scoring is labelled (red). (D-L) Estimation of hippocampal cell type absolute abundance by deconvolution of bulk RNAseq data from aging WT and TSPO-KO mice under inflammation using CIBERSORTx. Signature reference cell type markers were derived from single cell RNAseq of adult mouse hippocampus from (Zeisel *et al*., 2015). Units of the graphs are arbitrary. Data shown as median, interquartile range with error bars indicating minimum and maximum. Statistical tests: Fig 1A, B: Benjamini Hochberg (BH) corrected FDR <0.05; NS, non-significant. Fig 1C: Fisher’s Exact test to test. Fig 1D-L: Permutational univariate ANOVA, Dunn multiple comparison post-hoc test. \*\**p* < 0.01. * *p <* 0.05.

We directly tested the overlap between TSPO-KO specific transcriptional signatures in the young (Fairley *et al*., 2023) versus aged inflammatory hippocampus. Aging resulted in a reversal of TSPO-KO transcriptional signatures following inflammatory insult. A significant overlap between young *upregulated* and aged *downregulated* differentially expressed genes (DEG) (*p*<0.0001, odds ratio =11.39; Fig 1C, Supplementary Table 1C), and young *downregulated* and aged *upregulated* DEGs was observed in the inflammatory TSPO-KO hippocampus (*p*<0.0001, odds ratio =46.10; Fig 1C, Supplementary Table 1C). Functional annotation indicated genes that were upregulated in the young but downregulated in the aged inflammatory TSPO-KO hippocampus related to transmembrane and calcium ion transport and myelination. Meanwhile, genes that were downregulated in the young but upregulated in the aged inflammatory TSPO-KO hippocampus related to immune system processes, cellular response to interferons (β, γ) and regulation of TNF production. This indicated an interaction between TSPO and aging in inflammatory responses.

### 3.2 TSPO – aging interaction associated with changes in hippocampal cell heterogeneity

To determine if changes in cell composition, such as increased immune cell infiltration, may contribute to the TSPO-aging interaction in inflammatory transcriptional responses, we used computational deconvolution methods to estimate hippocampal cell-type composition from the bulk transcriptomic data in young versus aged WT and TSPO-KO mice treated with LPS. No changes in estimates of total populations of interneurons, pyramidal neurons, oligodendrocytes, astrocytes, macrophages or endothelial-mural cell populations were detected (Fig 1D-I, Supplementary Table 1D). However, an age-related decrease in estimates of a subtype of GABAergic interneurons (subtype 7) found in the hippocampal CA1 region, a glutamatergic pyramidal neuron subtype found in the subiculum (subiculum subtype), and a post-mitotic pre-myelinating oligodendrocyte subtype (subtype 1), were detected in LPS treated TSPO-KO but not WT hippocampus (Fig 1J; interneuron subtype 7: main effect: age x genotype, F = 28.44, *p* = 0.004; Dunn multiple comparison *p* = 0.009; pyramidal neuron subiculum subtype: age x genotype, F = 4.50, *p* = 0.05, Dunn post-hoc corrected for multiple comparisons young vs old TSPO-KO *p* = 0.02; oligodendrocyte subtype 1: age x genotype F = 8.53, *p =* 0.008, Dunn post-hoc corrected for multiple comparisons young vs aged TSPO-KO *p* = 0.045). This suggests that the downregulation of genes in pathways related to GABAergic synapse that were identified in aged TSPO-KO mice under inflammation (Fig 1B), may reflect neuronal death. Likewise, the age-related decrease in myelination-related genes observed in young versus aged TSPO-KO mice may be driven by reduced pre-myelinating oligodendrocytes.

With respect to the immune cells, there was a significant increase in the ratio of perivascular macrophages to microglia cell estimates in the hippocampus of the aged TSPO-KO mice under inflammation (main effect: age x genotype; F= 8.14, p = 0.02; Dunn pairwise post-hoc, corrected for multiple comparisons young TSPO-KO vs aged TSPO-KO *p* = 0.03). This change in macrophage composition reflected a greater proportion of perivascular macrophage populations relative to microglia. Absolute estimates of microglia revealed a decrease in a microglia subtype characterized by expression of *Hexb* in LPS treated, aged TSPO-KO mice (age x genotype F = 12.85, p = 0.002, Dunn post-hoc corrected for multiple comparisons p = 0.06; Fig 1K). While it did not reach significance after correction, this was surprising given that microglia are a key inflammatory mediator in the brain, and TSPO-KO was associated with severely exacerbated transcriptional inflammatory signals in aging. The gene signature of this microglial subtype was enriched for pathways involved in cytokine-cytokine receptor interaction, osteoclast differentiation and phagosome (Supplementary Table 1E). Likewise, absolute estimates of perivascular macrophages did not significantly change across treatments (age x genotype F = 16.15, *p* = 0.059, Supplementary Table 1F). However, interestingly, a proangiogenic subtype of perivascular macrophages (subtype 2) was detected exclusively in aged TSPO-KO mice under inflammation (Supplementary Table 1F). This was coupled with an increase in estimates of vascular endothelial cells subtype 1(age x genotype F= 28.99, *p*=0.002, Dunn posthoc multiple comparisons TSPO-KO aged vs young *p* = 0.02, WT aged vs TSPO-KO aged *p*=0.03), which was enriched with genes involved in cytokine-cytokine receptor interaction pathways (Fig 1L). Both the perivascular macrophage and the vascular endothelial subtype that were increased in the TSPO-KO in inflammaging were enriched in cytokine-cytokine receptor interaction pathways. Analysis of the Zeisel dataset (Zeisel *et al*, 2015) indicated that these perivascular and endothelial populations express high levels of TSPO in adult wild type hippocampus, even compared to microglia, although their relative estimated abundance was far lower than microglia. Upregulated TSPO expression in perivascular macrophages has been shown in a number of neuroinflammatory conditions including acute haemorrhagic leukoencephalopathy, viral infection and cerebrovascular disease (Nutma *et al*, 2021; Victorio *et al*, 2024). In the context of cerebrovascular disease, TSPO upregulation in perivascular macrophages occurred independently of microglia (Wright *et al*, 2020), suggesting that TSPO expression and function may potentially be regulated in an immune cell type and disease specific way. Future studies could address the cell specific immune functions of TSPO in these cell populations using conditional knockout mouse models.

### 3.3 TSPO - aging interaction linked to transcriptional control of interferon regulatory factors (IRFs)

To describe systems-level transcriptional variation and identify discrete gene co-expression patterns associated with aging and TSPO deletion we used multi weighted correlation network analysis (multiWGCNA). Thirteen co-expression modules were identified (Fig 2A, Supplementary Table 2A). To identify modules associated with aging and/or TSPO deletion, we then used factorial PERMANOVA analysis of multivariate gene expression for each module (Tommasini & Fogel, 2023). This approach enables module – multi-trait association for complex experimental design (main effects: age, genotype, age x genotype interaction). Six modules significantly differed across age and/or genotype (Modules 1-6; Supplementary Table 2B). Of these, five modules are enriched for functional terms using Gene Ontology over representation analysis (Module 1,3-6; Fig 2B-E; Supplementary Table 2B). Modules 3-5 correlated within an eigengene network, while modules 1 and 6 were isolated modules. This was reflected in the module-trait associations described below.

**Figure 2.**
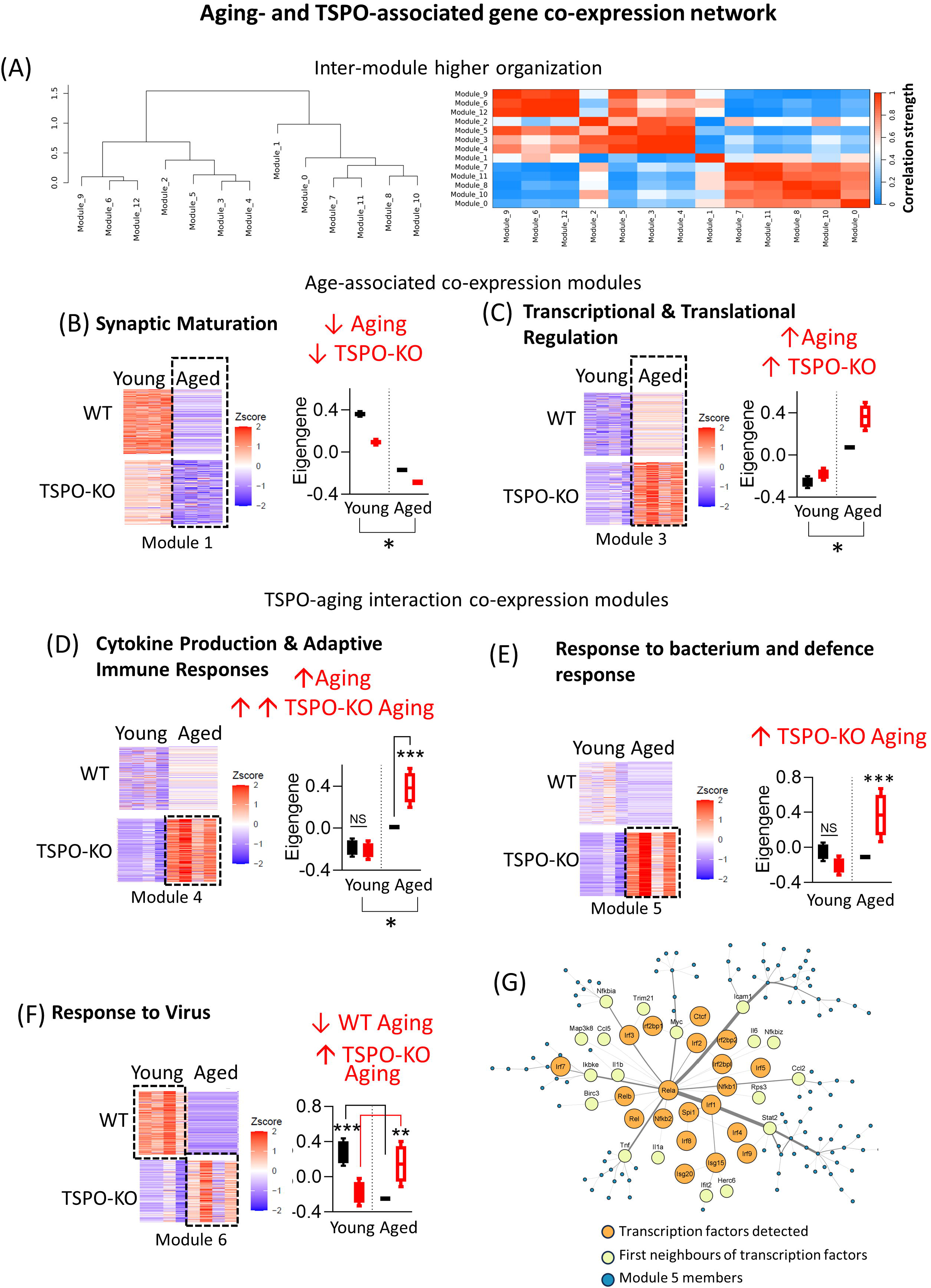
Multi WGCNA reveals co-expression network associated with TSPO in aging and inflammaging. (A) Cluster dendrogram and heatmap showing inter-module relationship among the modules detected. Representation of individual module is defined by the Module Eigengenes (first principal component, PC1, of each module). Hierarchical clustered dendrogram shows eigengene networks, which comprises of one/multiple module Eigengenes). Heatmap is calculated from the pairwise correlation between every module Eigengene, with intensity as strength of correlation. (B - E) Differential expression of modules functionally characterized by gene set enrichment analysis as (B) synaptic maturation module, (C) transcriptional and translational regulation module, (D) cytokine production & adaptive immune responses module, (E) response to bacterium and defense response module, and (F) response to virus module. For each module heatmap of expression of module members for young and aged WT and TSPO-KO samples shown (*left*) and graph of module expression summarized by the module eigengene for each group shown (*right*). Module expression, summarized by the module eigengene, or the PC1, is shown for each group. Eigengene data boxplots are inclusive of its median and interquartile range with error bars indicating minimum and maximum. (G) Integrated transcription factor – module members network of Module 5 (module exclusively upregulated by TSPO-KO Aged group). Edges refer to protein-protein interactions retrieved via STRING database. Orange nodes are the transcription factors whose motifs are significantly enriched (*BH p-adjusted value* <0.05) using Module 5 members as the input list to HOMER software. Yellow nodes refer to first neighbors of the transcription factors within Module 5. Transcription factors which have the highest degree connectivity (protein-protein interactions with the rest of module 5) *NF-Kβ* (e.g. *Rela*) and interferon transcription factors families (e.g *Irf1,Isg15*). Statistical tests: 2A-F: Multivariate PERMANOVA, EMM post-hoc test. * *p* < 0.05. ** *p* = 0.02. *** *p* < 0.002.

Modules 1 and 3 were age-associated co-expression modules. Module 1 was functionally associated with synaptic maturation; and was downregulated in both aging and TSPO-KO conditions (PERMANOVA: main effect age: *p =* 0.0001, genotype *p* = 0.0004, genotype x age *p* =0.0032; Supplementary Table 2A, B, Fig. 2B). This was consistent with previous aging studies showing age-associated downregulation of pathways related to synaptic maturation, plasticity and function in aging, which is thought to be a key factor in age-related cognitive decline (Smith *et al*, 2020). Interestingly, TSPO deletion exacerbated this age-associated depletion of synaptic transcriptomic signature in both young and aged mouse hippocampus.

Conversely, module 3 was functionally associated with immunity and transcriptional and translational regulation; and was upregulated in both aging and TSPO-KO conditions (PERMANOVA: main effect age: *p* = 0.004, main effect genotype: *p =* 0.0002; Fig. 2C). Immune-related members of module 3 included innate immunity and myeloid differentiation functional terms. These immune functions likely involving integrin-GTPase associate cell adhesion signalling processes, since *Itgb1* and *Cdc42* were identified as hub genes with the most protein-protein interactions network (PPI) constructed from the module driver genes members (Supplementary Figure 2A). Previous aging studies have also identified increased immune activation as a key signature in multiple regions of the aging brain (Ham & Lee, 2020). Meanwhile, ttranscriptional and translational regulatory functional terms in module 3 included genes involved in transcriptional elongation, regulation of RNA polymerase II promoter, and ribosome and ribonucleoprotein complex biogenesis (Supplementary Table 2B). Previous studies have shown age-related changes in transcriptional elongation and RNA splicing is associated with longevity across multiple organisms across different tissues including brain and mutations that alter RNA polymerase II activity increase longevity (Debès *et al*, 2023). Since these immune and transcriptional genes were identified as part of the same coregulated network, one possible explanation is these age-relate changes in immunity and transcription/translation are causally linked. Supporting this notion, several potential module driver genes identified by high intra-modular connectivity were found to act as dual members of immune and transcriptional regulatory functional terms. These genes included *Lpxn, Zfp36, Lgals3, Elf1, Cd33*. Similar to observations in the synaptic maturation module, TSPO deletion exacerbated these age-related transcriptomic signatures of module 3 in both young and aged mouse hippocampus. These findings indicate TSPO deletion exacerbates age-related transcriptional changes in the hippocampus.

Modules 4-6 were associated with the TSPO-aging interaction. Module 4 was identified as an immune module, associated with cytokine production and adaptive immune responses, and like module 3, was also upregulated in aging (PERMANOVA: main effect age: *p =* 0.0004; Supplementary Table 2B). Supporting the functional enrichment results, a PPI subnetwork of immune related transcription factors and helicases (*Stat3-Jak1/2-Rela*) were enriched using the STRING database (Supplementary Figure 2C). Interestingly, this age-related increase in module 4 was markedly exacerbated in the TSPO-KO condition (age x genotype: *p* = 0.0045, EMM post-hoc aged WT s aged TSPO-KO *p* = 0.00013; Fig. 2D). Meanwhile another immune module (module 6), involved in viral responses and innate immunity, was downregulated in aging in WT, but upregulated in aging in TSPO-KO hippocampus (PERMANOVA: age x genotype p= 0.0001; EMM post-hoc: WT young vs old *p =* 0.001, TSPO-KO young vs old *p =* 0.018; Fig. 2E, Supplementary Table 2B). Previous studies have demonstrated TSPO-PET can be used to visualize immune responses to viral infections (Shah *et al*, 2022). Our data suggests TSPO may play a role in dampening viral immune responses in aging hippocampus, and that this TSPO sensitivity to viral infections may be regulated by receptors involved in antigen presentation, given that the top 5 members with highest intramodulatory connectivity were *Klhl21, Clec4a3, Gbp2, Trim30a, Iigp1* (Supplementary Table 2C).

An immune module that was functionally associated with response to bacterium and regulation of defense response (module 5) was of the highest interest amongst the coregulatory network because it was exclusively upregulated within the aged TSPO-KO group (PERMANOVA: age x genotype *p* = 0.0001; TSPO-Fig. 2D, EMM posthoc aged TSPO-KO vs all other groups *p* < 0.002; Supplementary Table 2B). To predict potential transcription factors involved in regulating this TSPO-aging interaction, we leveraged motif detections of promoters (-500bp to +2000bp from TSS) based on the members of this module (module 5; Fig 2G). Nine motifs of transcription factors were significantly enriched (FDR <0.05; Supplementary Table 2D). A total of 20 cognate genes (transcription factors) of these motifs were found within the expression matrix. These were integrated as a protein-protein interaction network with module 5 members (Fig 2G). *Rela*, a transcription factor within the Nuclear Factor Kappa B (*NF-kβ)* family (Yu *et al*, 2004), was identified as the hub node based on the highest degree connectivity (21 protein-protein interactions) and shortest average path (3.027) within the network (Supplementary table 2D). Further, most of the mapped transcription factors (13/20) belonged to interferon transcription factors family (e.g *Irf1,Isg15*). Interestingly, *Spi1*, a macrophage-microglia transcription factor (Zhang *et al*, 2024), is predicted to interact with enriched interferon transcription factors (*Irf1,Irf4,Irf8*), suggesting that macrophage and/or microglia may be involved in mediating the interferon signalling network seen in module 5 (Fig 2G). Together, the integrated network of transcription factor-module 5 indicates that the immune pathways enriched within module 5 may be largely mediated by *NF-kβ* and interferon signalling. Our finding indicating the link between TSPO and *NF-kβ* is not an isolated finding. While a previous study found that TSPO induces a retrograde mitochondrial-nuclear signalling via *NF-kβ* in breast cancer cell line survival (Desai *et al*), our data suggests the age-dependent effects of TSPO on immune responses may be mediated via an effect on *NF-kβ* and interferon transcriptional pathways. Since *NF-kβ* and interferons have been identified as potential therapeutic targets in AD (Chavoshinezhad *et al*, 2023; Grimaldi *et al*, 2014; Mudò *et al*, 2019; Sun *et al*, 2022), this TSPO-aging interaction in *NF-kβ* and interferon transcriptional pathways may be an important consideration in therapeutic development.

### 3.4 Drugs that disrupt cell cycle and induce DNA-damage mimic TSPO-dependent aging transcriptional signature

To identify small molecules that phenocopy TSPO-dependent inflammaging, we compared the TSPO-dependent inflammaging transcriptome signature with drug gene expression signatures in a perturbational signature library called Connectivity Map (CMap), using the LINCS database. The similarity (or dissimilarity) between the TSPO-KO inflammaging and drug signatures is quantified with a connectivity score, where a score of +1 indicates a strong transcriptional phenocopy, while a score of –1 indicates a strong transcriptional reversal of the TSPO-KO inflammaging signature. We focused on transcriptional signatures from drugs screened in two key cell types: neural progenitor cells (NPCs) and a glioblastoma cell line (GI1).

CDK (cyclin-dependent kinase) inhibitors, topoisomerase inhibitors and heat shock protein 90 (HSP90) inhibitors were commonly identified in both cell types analysed to closely mimic the inflammatory transcriptional signature characterizing TSPO-dependent aging (Fig 3A, B) The highest ranked phenocopy compound identified in the NSCs, Dinaciclib (mean connectivity score = +0.33), is a selective inhibitor of the CDK1, CDK2, CDK5 and CKD9. This compound was not tested in the GI1 cell line. CDK inhibitors suppress the activity of key kinases at different stages of the cell cycle process thus halting cell proliferation (Parry *et al*, 2010; Webster & Kimball, 2000). Another selective CDK1 inhibitor, CGP-60474, is the 5th ranked phenocopying compound with TSPO-KO inflammaging (mean connectivity score = +0.30; Fig 3A). Likewise, a number of CDK inhibitors were also identified to significantly phenocopy the TSPO-dependent inflammaging signature in the glioma GI1 cells, although they were not the strongest phenocopies identified (Supplementary Table 3). Although specific CDK inhibitors were not as strongly correlated with the TSPO-KO inflammaging signature in the GI1 glioma compared to the NPC, the top ranked phenocopy drug in the GI1 glioma cells is the CHK (Checkpoint) inhibitor, LY-2606368 (1st, +0.25). Checkpoint kinases are upstream regulators of CDKs (Janetka & Ashwell, 2009), corroborating potential disruption of cell cycle processes in TSPO-KO inflammaging.

**Figure 3.**
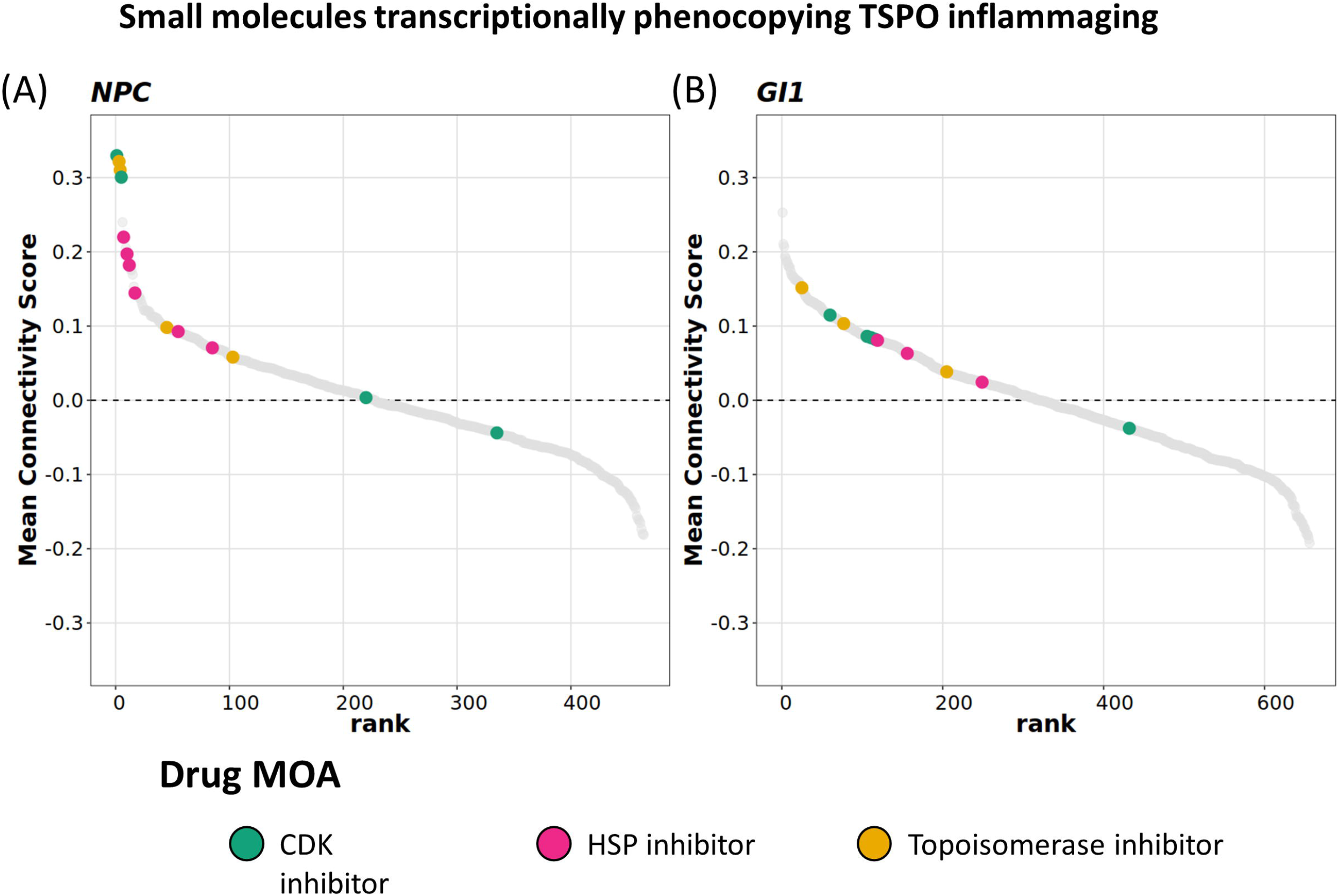
Small molecules identified to transcriptionally phenocopy the TSPO inflammaging signature via Connectivity Mapping. Mean connectivity score against rank of LINCS compounds treated in (A) NPC and (B) GI1 cell line when queried with TSPO inflammaging signature. Positive connectivity score indicates the phenocopying effect of a compound with the query signature, while a negative score indicates the anti-correlation between the compound and query signature. Compounds are ranked in ascending order, from the strongest phenocopy to strongest reversal signal. Green: CDK inhibitors, Pink: HSP90 inhibitors, Yellow: Topoisomerase inhibitors.

Two Topoisomerase inhibitors, Mitoxantrone and Topotecan, also strongly phenocopied the TSPO-KO inflammaging signature in NSCs (ranked 2nd and 3rd, mean connectivity score = +0.32 and +0.31, respectively; Fig 3A). Although these two topoisomerase inhibitors were not tested in the GI1 glioma cell line, the inhibitors Daunorubicin (25th rank, mean connectivity score = +0.15) and Camptothecin (77th, +0.10) also significantly phenocopied the TSPO-dependent inflammaging signature (Fig. 3B). Topoisomerase inhibitors are DNA-damage inducing compound, that act by trapping the Top1-DNA cleavage complex (Top1cc) and preventing the cleaved DNA from re-ligating, typically used to trigger cancer cell apoptosis (Katyal *et al*, 2014). This potentially indicated elevated DNA-damage in TSPO-KO inflammaging hippocampus.

Interestingly, two heat shock protein 90 (HSP90) inhibitors were also highly ranked, phenocopying the TSPO-KO inflammaging signature — Tanespimycin (ranked 7th, mean connectivity score = +0.22) and Geldanamycin (ranked 10th, mean connectivity score = +0.20; Fig 3A). These compounds bind to the HSP90, which in turn releases the HSF1 (heat shock factor 1) transcription factor to activate the expression of other HSPs during cellular stress (Kurop *et al*, 2021). This finding suggests that cellular stress triggering the heat shock response may also be associated with the loss of TSPO function during inflammaging.

### 3.5 TSPO deletion disrupts correlation of AD risk genes in aging hippocampus under inflammation

Since TSPO expression is increased in AD brains and we have previously shown TSPO deletion affects AD-related pathogenesis in mice, we investigated the effect of TSPO deletion on expression of AD risk genes. Among the mouse orthologs of 20 AD risk genes, *Tspo* expression significantly correlated with the expression of 13 AD risk genes in the hippocampus of both young and aged WT mice under inflammation (FDR <0.05). *Tspo* expression positively correlated with expression of *Ap2a2, Apoc1, Frmd4a*, *Gab2*, *Mthfd1l*, *Ptk2b*, *Rin3* and *Sqstm1;* and negatively correlated with *Bin1, Mef2c, Siglech, Sorl1* and *Trem2* expression (Fig. 4A; Supplementary table 4). The strength of correlation increased in aging between *Tspo* and the AD risk genes: *Rin3, Sqstm1, Mthfd1, Frmd4a, Trem2, Siglech,* and *Bin1.* In the aged but not young hippocampus, *Tspo* expression additionally strongly correlated with expression of *CD33* and *Apoe* and negatively with *Sppl2a.* Meanwhile, *Inpp5d* correlated with TSPO expression in young but not aged hippocampus (Supplementary table 4). Of the 13 AD risk genes that significantly correlated with *Tspo* expression, 6 were immune–associated genes (*Cd33, Gab2, Inpp5d, Mef2c, Ptk2b, Trem2 and Siglech),* the majority of which were age-dependently differentially expressed in the inflamed hippocampus of TSPO-KO but not WT mice (young vs aged TSPO-KO, upregulated: *Cd33, Gab2, Inpp5d, Ptk2b;* downregulated *Mef2c* and *Siglech,* Supplementary table 4). Other AD risk genes that do not exhibit cell-specific expression patterns including *Mthfd1l* and *Rin3* were also age-dependently differentially expressed in inflamed TSPO-KO hippocampus.

**Figure 4.**
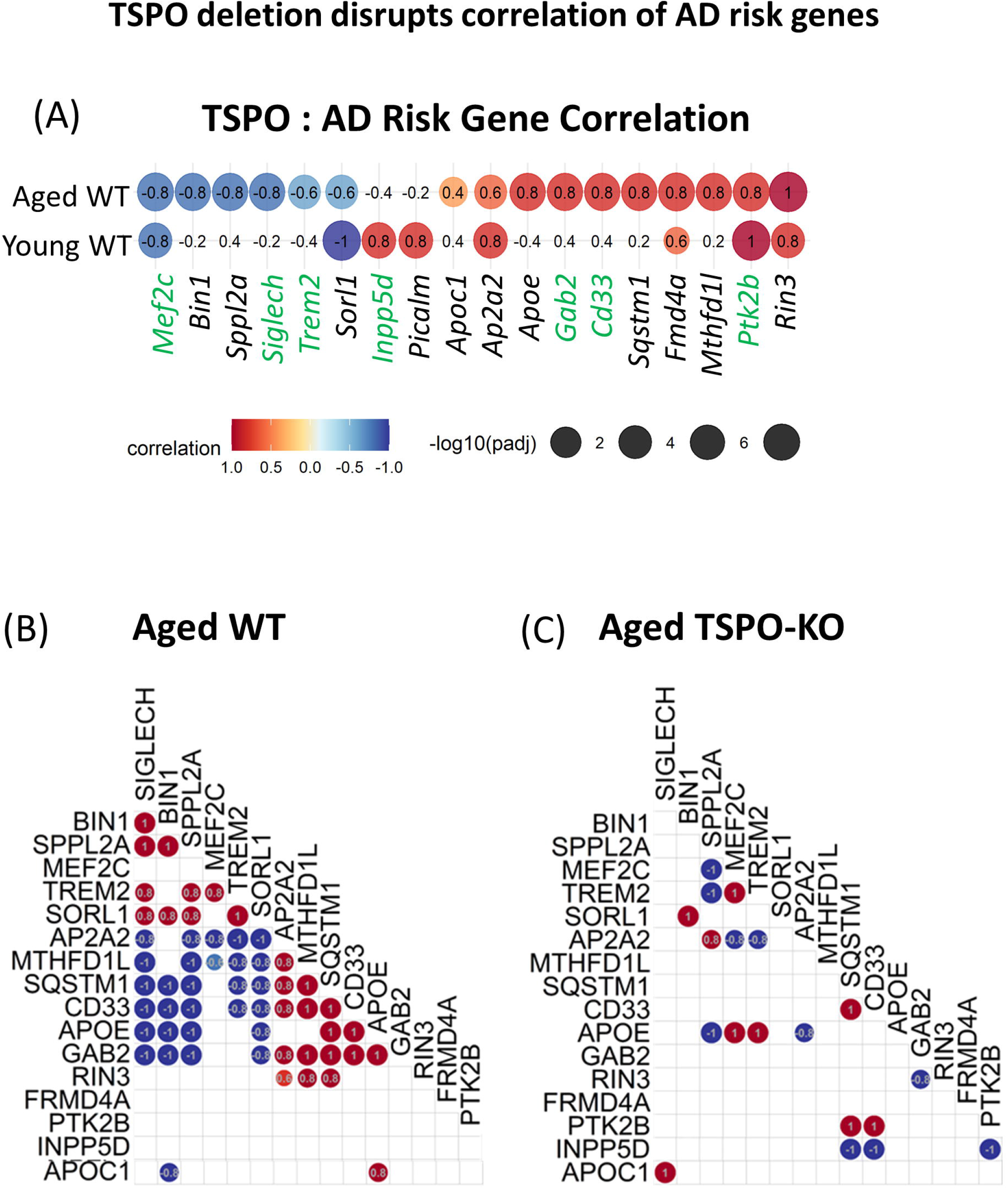
TSPO deletion disrupts correlation of AD risk genes in aging hippocampus under inflammation. (A) Correlation between transcriptional expression of *Tspo* and mouse orthologs of AD risk genes in young (3mo old) and aged (20 mo old) WT mouse hippocampus. Immune-related AD risk genes indicated in green font. Strength of correlation indicated by R^2^ value. For significant correlations (*p* < 0.05) R^2^ shown as dotplot heatmap, with significance indicated by dot size. (B, C) Significant correlation between transcriptional expression of AD risk genes in aged WT hippocampus (B) and aged TSPO-KO hippocampus (C). Strength of correlation measured by R^2^ shown as heatmap.

Correlation analysis between expression of AD risk genes in the hippocampus revealed differences in the aged WT and TSPO-KO hippocampus under inflammation (Fig 4B-–C). In the aged WT hippocampus, a high degree of correlation was observed between AD risk genes (Fig. 4B), which was disrupted in the TSPO-KO mouse hippocampus (Fig. 4C). In normal aging hippocampus, expression of the microglial receptor, *Cd33,* strongly negatively correlated with its downstream inhibitor, *Trem2* (Griciuc *et al*, 2019) and *Siglech,* which is a marker of homeostatic phagocytic microglia (Yao *et al*, 2022). This was not observed in aged TSPO-KO hippocampus. Instead, in the aged TSPO-KO hippocampus *Trem2* strongly correlated with expression of *Apoe* and *Mef2c.* Activation of APOE-TREM2 signalling, was previously found to be associated with neurodegenerative microglia phenotype (Krasemann *et al*, 2017).

### 3.6 Aging and TSPO alter brain metabolic profiles in inflammation

Since our transcriptomic data identified TSPO-dependent changes in metabolic pathways in the aged brain under inflammation, NMR was used to determine brain metabolites involved in metabolism of carbohydrates, amino acids and lipids in young and aged WT and TSPO-KO mice treated with LPS. We identified 19 metabolites which includes amino acids (glutamine, glutamate, aspartate, N-acetylaspartate (NAA), gamma-aminobutryric acid (GABA), tricarboxylic acid cycle (TCA) metabolites (fumarate, succinate), redox cofactors (NAD, Nicotinamide (NAM), Adenosine diphosphate (ADP) and membrane components, (choline, glycerophosphatidylcholine, Phosphatidylcholine) (Fig 5A). Additionally, a peak corresponding to formaldehyde, which is usually undetectable in brain, was identified exclusively in the aged TSPO-KO mice (Fig 5B).

**Figure 5.**
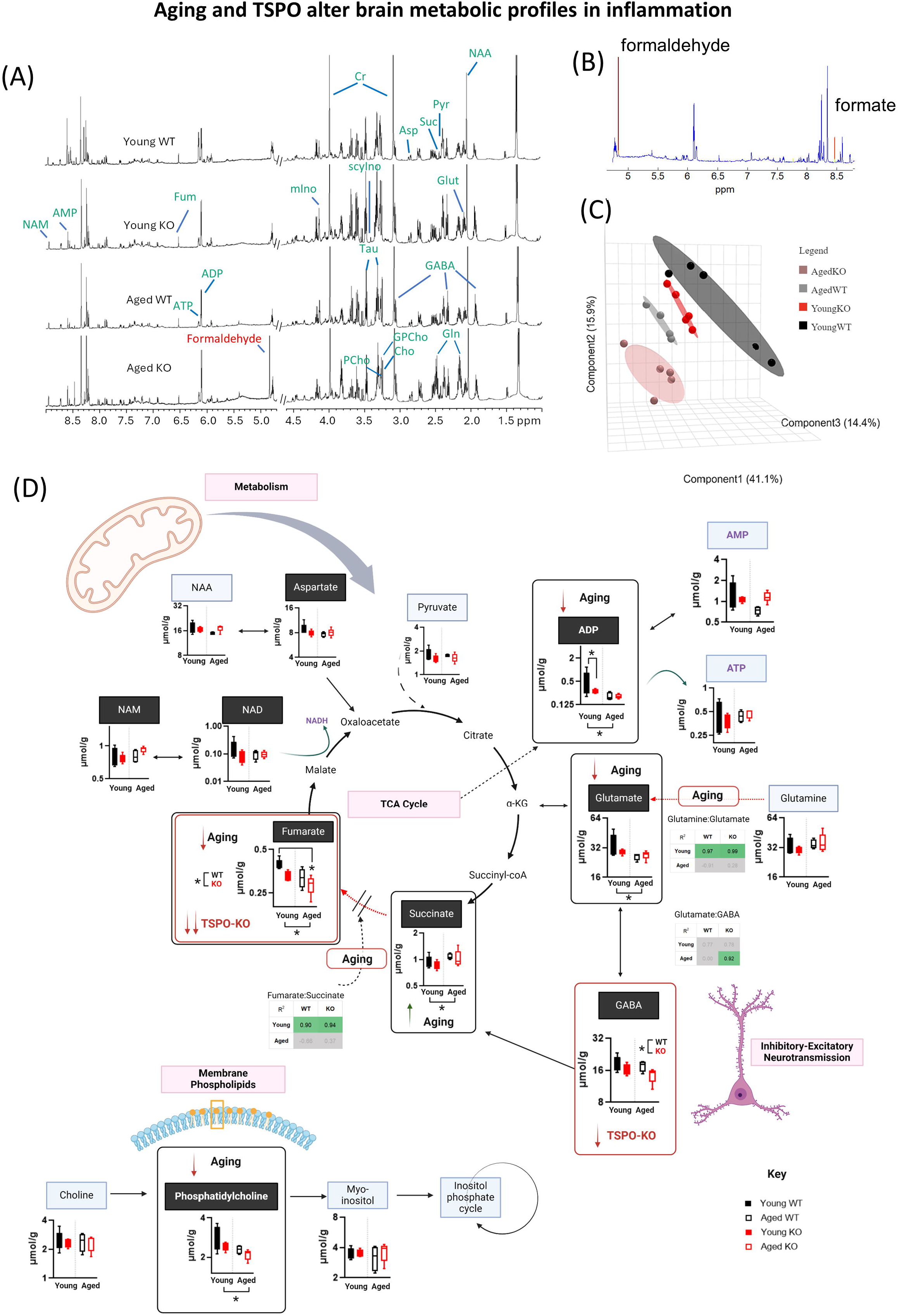
Aging and TSPO alter brain metabolic profiles in inflammation. (A) Representative 1D-NOSEY spectra of the 19 metabolites detected in young WT, old WT, young KO and old KO brain tissue samples. Metabolite assignment to NMR peaks indicated. Abbreviation: NAA, N-acetyl aspartate; gln, glutamine; glu, glutamate; Suc, succinate; Tau, taurine; scyIno, scyllo-inositol; mIno, myo-inositol; GPCho, glycerophosphocholine; PCho, phosphocholine; Cho, choline; Cr, creatine; GABA, γ-aminobutyrate; Pyr, pyruvate; Fum, fumarate; AMP, Adenosine monophosphate; ADP, Adenosine diphosphate; ATP, Adenosine triphosphate; Nam, nicotinamide. (B) STOCSY shows high correlation (0.8) between the two singlets (red peaks) at 4.832 (identified in aged TSPO-KO only) and 8.462 ppm, which is formic acid. The correlation suggests that peak at 4.832 could be formaldehyde. Formaldehyde in water presents structure of methanediol and it is a singlet with chemical shift ranging from 4.4-5.4 ppm (Automated Topology Builder (ATB) and Repository Version 3.0, Methanediol | CH4O2 | MD Topology | NMR | X-Ray (uq.edu.au)). (C) Multivariate Dimensionality reduction of targeted NMR metabolites using Partial least squares-discriminant analysis (PLSDA) for young and aged WT and TSPO-KO mice under LPS-induced inflammation. (D) Quantification of metabolites measured by NMR in young and aged brain of WT and TSPO-KO mice under LPS-induced inflammation. Metabolite concentrations shown as median, interquartile range with error bars indicating minimum and maximum. Metabolites that contributed the greatest variance to discrimination of the groups along PC1 are highlighted with black header (aspartate, NAM, NAD, ADP, glutamate, GABA, succinate, fumarate and phosphatidylcholine). Correlation heatmaps show Precursor:Product ratios as measured by Pearsons correlation, with significance threshold *p*<0.05. Precursor:Product metabolite ratios which showed significant changes during aging were Fumarate:Succinate, and Glutamine:Glutamate (Fumarate:Succinate and Glutamine:Glutamate significantly correlated in the Young but not Aged). Conversely, Glutamate:GABA ratio showed significant correlation exclusively in the TSPO Aged group.

PLS-DA was used to determine the major metabolites discriminating between young versus aged and WT versus TSPO-KO conditions (Fig 5C). Each group clustered with distinct separation by genotype for each age. Aging was most clearly discriminated along the principal component axis 1 and 2, which explained 41.1% and 15.9% of the variability, respectively (Fig 5C). Metabolites that contributed the greatest variance to discrimination of the groups along PC1 were involved in the TCA cycle (Fumarate, Succinate, Glutamate, GABA, Aspartate), membrane phospholipids (Phosphatidylcholine, Glycerolphosphocholine), and redox cofactors (NAD, NAM, ADP; Variance Importance Projections (VIP) score > 0.8; Fig 5A).

Comparison of metabolite concentrations in aging revealed increased levels of the TCA metabolite, succinate, coupled with reduced levels of its product, fumarate, suggesting inefficient conversion of succinate to fumarate in the aged inflammatory brain (PERMANOVA: main effect: aging; Fumarate, F_(1,5.66)_ = 8.41, *p* = 0.007; succinate, F_(1,5.66)_ = 4.98, *p* = 0.043; Fig 5D, Supplementary Table 5). Supporting this, succinate and fumarate levels were significantly correlated in young but not aged brains in both WT and TSPO-KO mice, suggesting a break in the TCA cycle at succinate dehydrogenase in aged mice (Fig. 5D). TSPO deletion was also associated with depleted levels of fumarate (PERMANOVA: main effect: genotype; Fumarate, F_(1,5.66)_ = 5.82, *p* = 0.026; Fig 5D, Supplementary Table 5). Further, age-related depletion of fumarate was more severe in TSPO-KO brains, with reduced levels detected in young and aged TSPO-KO mice compared to young WT (PERMANOVA: age x genotype interaction; Supplementary Table 5; fumarate, F_(3,5.66)_ = 6.70, *p* = 0.003; young WT vs aged TSPO-KO, FDR = 0.0067). Interestingly, fumarate is an anti-inflammatory metabolite, shown to inhibit activation of NF-κβ and approved for treatment of multiple sclerosis (Miljković *et al*, 2015). Therefore, depleted levels of this metabolite could potentially contribute to the increased inflammatory transcriptional signatures detected in TSPO-KO inflammaging.

Age-related reductions in levels of glutamate, which fuels the TCA cycle via α-ketoglutarate, in addition to being the major excitatory neurotransmitter of the brain, were also observed (PERMANOVA: main effect: aging; glutamate, F_(1,5.66)_ = 5.70, *p* = 0.011; Fig 5D, Supplementary Table 5). Glutamate concentrations significantly correlated with levels of its precursor, glutamine in young but not aged brains of both WT and TSPO-KO mice, potentially indicating reduced efficiency of glutamate synthesis in the aging brain (Fig. 5D). In contrast, TSPO deletion resulted in significantly reduced levels of the glutamate metabolite, GABA, which feeds into the TCA cycle via succinate, in addition to being the major inhibitory neurotransmitter of the brain (PERMANOVA: main effect: genotype; GABA, F_(1,5.66)_ = 6.02, *p* = 0.015; Fig 5D, Supplementary Table 5). Depleted brain GABA levels in TSPO-KO mice was consistent with our transcriptomic findings, that identified a downregulation of pathways related to the GABAergic synapse in the hippocampus of aged TSPO-KO vs WT mice. These changes in glutamate and GABA levels could potentially reflect excitatory-inhibitory imbalance, which is associated with anxiety and depression(Zhang *et al*, 2021). We have previously shown that TSPO deletion increases anxiety-related behavior in mice (Barron *et al*, 2021), thus changes in bioavailability of glutamate and GABA may contribute to this effect.

Reduced accumulation of the energy substrate, ADP, was also observed in the aging brain (PERMANOVA: main effect: aging; ADP, F_(1,5.66)_ = 4.65, *p* = 0.005; Fig. 5D). ADP levels significantly correlated with concentrations of its metabolite, AMP, but not ATP, in the brains of young WT mice only. A significant interaction between TSPO-KO and aging was identified for ADP, with pairwise analysis indicating ADP was significantly reduced due to aging in WT mice only (PERMANOVA: age x genotype interaction; Supplementary table 5; ADP, F_(1,5.66)_ = 2.60, *p* = 0.026; Fig 5D).

Lastly, reduced levels of the membrane phospholipid, phosphatidylcholine, were also observed in aging brain in inflammation (PERMANOVA: main effect: aging; phosphatidylcholine, F_(1,5.66)_ = 6.77, *p* = 0.013; (Fig 5D, Supplementary Table 5). In young mice but not aged mice, phosphatidylcholine concentrations positively correlated with glycerophosphatidylcholine concentrations (WT young r^2^ = 0.89, *p* = 0.04; TSPO-KO young r^2^ = 0.93, *p* = 0.02; WT old r^2^ = -0.79, *p* = 0.20; TSPO-KO young r^2^ = 0.056, *p* = 0.92). A significant interaction between TSPO-KO and aging was also identified for phosphatidylcholine (PERMANOVA: age x genotype interaction; Supplementary Table 5; phosphatidylcholine, F_(1,5.66)_ = 3.63, *p* = 0.035 (Fig 5D). Previous studies have demonstrated hippocampal phospholipid levels including phosphatidylcholine decline in aging. In addition to comprising the cell membrane, phosphatidylcholine is also a precursor for the major neurotransmitter acetylcholine(Chung *et al*, 1995), which is important in memory and depleted in AD (Giacobini *et al*, 2022; Liu *et al*, 2022). Interestingly, phosphatidylcholine has bene shown to inhibit LPS-induced inflammation and improve cognitive function (Tan *et al*, 2020), opening the possibility that age-related decline in hippocampal phosphatidylcholine may also play a role in inflammaging.

## 4 Conclusions

Broadly, our study indicates TSPO plays a protective role in brain aging, and that TSPO deletion exacerbates age-related transcriptional and metabolic changes in the hippocampus. TSPO deletion aggravated key age-related synaptic and immune transcriptional signatures in the inflamed hippocampus, and these inflammaging signatures were mimicked by drugs that disrupt the cell cycle, cause DNA-damage and cell stress through the heat shock response. TSPO deletion also worsened age-related changes in brain metabolites, including depletion of the major inhibitory neurotransmitter GABA. Importantly, we found an interaction between TSPO-function and aging in the hippocampus. Aging resulted in a reversal of TSPO-KO transcriptional signatures following inflammatory insult. While we have previously shown that TSPO deletion dampens hippocampal inflammatory signatures in the young adult, we were surprised to discover that loss of TSPO drastically exacerbated inflammatory transcriptional responses in the aging hippocampus. This TSPO-aging interaction was linked to transcriptional control of interferon regulatory factors. Interferons have been implicated in brain aging as well as the pathogenesis of AD. Interestingly, while expression of AD risk genes were highly correlated in the normal aging hippocampus, this was disrupted in the absence of TSPO, supporting the notion that TSPO regulates molecular pathways implicated in AD pathogenesis. This TSPO-aging interaction is an important consideration in the interpretation of TSPO-targeted biomarker and therapeutic studies, as well as *in vitro* studies which cannot model the aging brain.

## CRediT authorship contribution statement

**Kei Onn Lai:** Conceptualization, Methodology, Investigation, Validation, Formal Analysis, Visualization, Writing – original draft. **Nevin Tham:** Methodology, Investigation, Validation, Formal Analysis, Visualization, Writing – original draft. **Lauren Fairley:** Methodology, Investigation, **Roshan Naik:** Methodology, Investigation. **Yulan Wang:** Methodology, Investigation, Validation, Formal Analysis, Visualization, Writing – review & editing. **Sarah R. Langley:** Methodology, Writing – review & editing. **Anna M. Barron:** Conceptualization, Formal Analysis, Visualization, Writing - original draft, Supervision, Funding acquisition.

## Acknowledgements

This project was funded by the Nanyang Assistant Professorship Award from Nanyang Technological University Singapore (AMB).

## Data availability

Study data will be made available via https://researchdata.ntu.edu.sg/dataverse/neurobiologyagingrepository.

