## Supplementary Figure 2 for "Interaction between mitochondrial translocator protein and aging in inflammatory responses in mouse hippocampus"

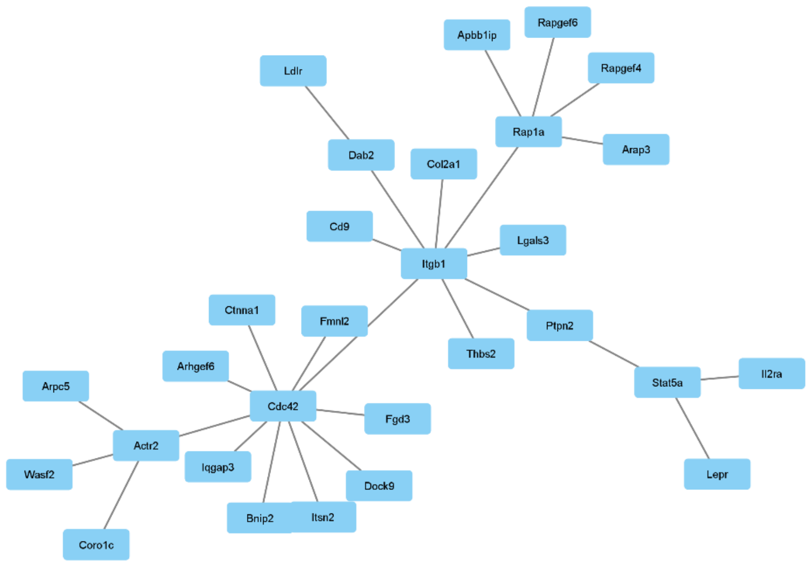


Supplementary Figure 2A: Immune subnetwork of PPI interactions derived from the module driver genes of Module 3. Network edges refer to direct PPI interactions queried from STRING database via NetworkAnalyst.

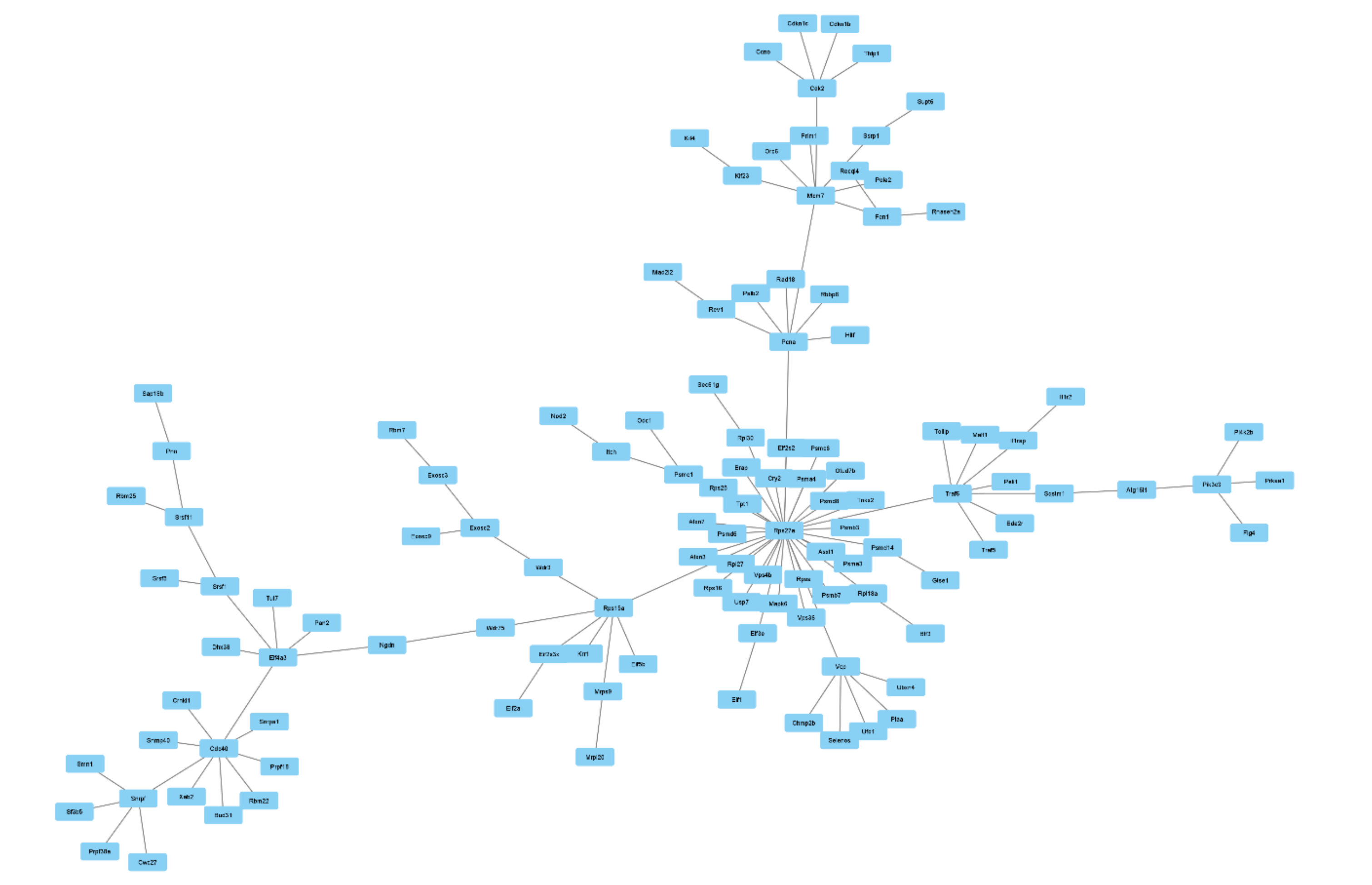

Supplementary Figure 2B:Transcriptional-Translational processes subnetwork of PPI interactions derived from the module driver genes of Module 3. Network edges refer to direct PPI interactions queried from STRING database via NetworkAnalyst.

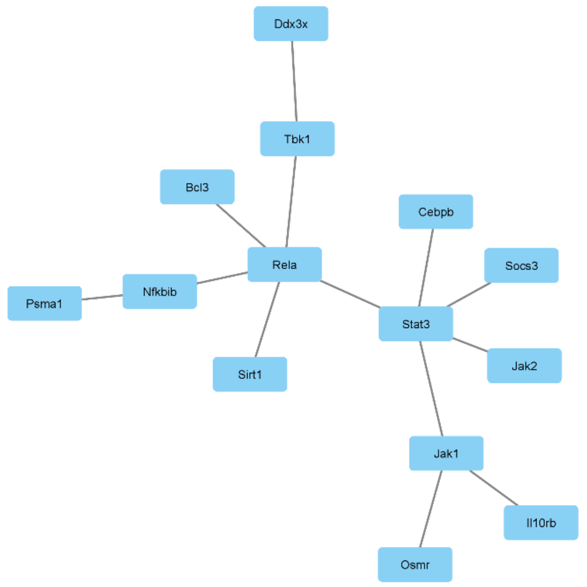


Supplementary Figure 2C: PPI subnetwork of immune related transcription factors and helicases derived from the module driver genes of Module 4. Network edges refer to direct PPI interactions queried from STRING database via NetworkAnalyst.
